# Systemic CD8+ T cell effector signature predicts prognosis of lung cancer immunotherapy

**DOI:** 10.1101/2024.09.16.613381

**Authors:** Hyungtai Sim, Geun-Ho Park, Woong-Yang Park, Se-Hoon Lee, Murim Choi

## Abstract

**Background:** While immune checkpoint inhibitors (ICIs) are adopted as standard therapy in non-small cell lung cancer (NSCLC) patients, factors that influence variable prognosis still remain elusive. Therefore, a deeper understanding is needed of how germline variants regulate the transcriptomes of circulating immune cells in metastasis, and ultimately influence immunotherapy outcomes.

**Methods:** We collected peripheral blood mononuclear cells (PBMCs) from 73 ICI-treated NSCLC patients, conducted single-cell RNA sequencing, and called germline variants via SNP microarray. Determination of expression quantitative trait loci (eQTL) allows elucidating genetic interactions between germline variants and gene expression. Utilizing aggregation-based eQTL mapping and network analysis across eight blood cell types, we sought cell-type-specific and ICI-prognosis-dependent gene regulatory signatures.

**Results:** Our sc-eQTL analysis identified 3,616 blood- and 702 lung-cancer-specific eGenes across eight major clusters and treatment conditions, highlighting involvement of immune-related pathways. Network analysis revealed TBX21-EOMES regulons activity in CD8+ T cells and the enrichment of eQTLs in higher-centrality genes as predictive factors of ICI response.

**Conclusions:** Our findings suggest that in the circulating immune cells of NSCLC patients, transcriptomic regulation differs in a cell type- and treatment-specific manner. They further highlight the role of eQTL loci as broad controllers of ICI-prognosis-predicting gene networks. The predictive networks and identification of eQTL contributions can lead to deeper understanding and personalized ICI therapy response prediction based on germline variants.

## Background

Lung cancer is the leading cause of cancer-related death worldwide [1]. While smoking is the predominant risk factor, germline variants also contribute to disease susceptibility, with a genotype-based heritability of 10–20% [2]. Large-scale genome-wide association studies (GWAS) have identified over a thousand significant loci associated with lung cancer, implicating not only genes involved in smoking behaviors and DNA repair processes, but also genes with uncharacterized functions [3].

During tumor pathogenesis, practically all hematopoietic lineages undergo phenotypic and proportional changes in systemic immunity [4]. For instance, immunosuppressive cell types, such as regulatory T cells and immature neutrophiles, are expanded in cancer patients [5, 6]. Additionally, impaired activation of natural killer (NK) cells through transcriptional inhibition of the *NCR3* gene has been observed in non-small cell lung cancer (NSCLC), and is associated with a negative prognosis [7].

Immune checkpoint inhibitors (ICIs) have revolutionized the treatment of advanced NSCLC but with considerable variability in patient response and prognosis [8]. PD-L1 expression in tumors has been identified as a key biomarker for ICI treatment result, but recent studies emphasize a crucial role of systemic immunity in ICI efficacy. For example, systemic chemotherapy can negatively impact ICI efficacy by disrupting the immune system, whereas locally focused chemotherapy enhances ICI efficacy [9].

Moreover, specialized dendritic cells have been demonstrated crucial for presenting tumor-specific antigens to draining lymph nodes, thereby priming CD8+ T cells in a systemic manner [10]. While these findings represent promising progress, the genetic features of NSCLC patients that determine response to ICI remain to be discovered. Expression quantitative trait loci (eQTL) studies have emerged as a prominent tool for fine-mapping GWAS results by providing insights into how genetic variants influence gene expression across various perturbations, including disease status and environmental stresses [11, 12]. Single-cell RNA sequencing (scRNA-seq) enhances this approach by enabling the dissection of transcriptomic changes at a single-cell resolution, which facilitates the study of cell-type-specific and condition-specific gene expression regulation [13, 14]. In lung cancer, eQTL mapping has previously been conducted based on gene expression data from lung tissues [15–18] and by leveraging eQTL data from resources such as the Genotype-Tissue Expression (GTEx) project to identify colocalized GWAS results [19, 20].

Understanding the genetic mechanisms that underlie regulation of systemic immune responses in the context of metastatic tumors could provide new insights into cancer treatment, by improving prediction of ICI efficacy. Here, we leveraged single-cell eQTL (sc-eQTL) analysis to explore differences in the gene regulatory landscapes of NSCLC patients treated with ICIs, focusing on germline variants. By analyzing peripheral blood mononuclear cells (PBMCs) from 73 ICI-treated donors using scRNA-seq and SNP microarray, we aimed to uncover cell-type-specific and ICI-treatment-dependent gene regulatory signatures. Our findings indicate that eQTLs are enriched within genes having high centrality in the transcriptomic networks of ICI-treated patients, suggesting that germline variations have broader impacts upon treatment response in advanced NSCLC.

## Methods

### Patients

The cohort used in the study was previously described [27, 28]. Briefly, 73 stage IV NSCLC patients (36 LUAD, 30 LUSC, and 7 others) treated with anti-PD(L)1 (e.g., Atezolizumab, Nivolumab, and Pembrolizumab) were recruited at Samsung Medical Center (SMC) under the permission of the Institutional Review Board (No. 2018-04-048). Whole blood samples were collected before and 1-3 weeks after the treatment, PBMCs were isolated, and scRNA-seq and SNP microarray experiments were conducted. Responses to ICIs were clinically determined and categorized as either partial response (PR), progressive disease (PD), or stable disease (SD) (Additional file 1: Table S1). In the subsequent analyses, we further binarized responses from PR or PD patients according to progression-free survival (PFS) > 6 months. The participants did not carry significant loss-of-function variant in genes involved in immune checkpoint, interferon signaling, and DNA damage repair pathways (data not shown).

### Single-cell RNA-seq (scRNA-seq) data processing

Single-cell transcriptome data were generated using the Chromium single cell platform (10x Genomics, Pleasanton, CA) as previously described [27]. Raw fastq files were mapped using Cellranger version 7.0.1 on provided GRCh38 reference. Potential contaminated cell barcodes were filtered using cellbender and the identity of each cell was determined from imputed genotypes using demuxlet [29, 30]. Seurat v5 was used to preprocess raw count matrices, in which cells having mitochondrial reads <20%, and 200-6,000 expressed genes were retained [31]. We calculated SCTransform-normalized counts across each sample and corrected them by median counts using PrepSCTfindmarkers [22]. Based on the normalized count, scRNA-seq samples were integrated using Harmony and cell clusters defined by the Leiden algorithm [32]. Cluster annotations were validated based on marker genes and azimuth reference [31]. Finally, the UMAP algorithm was used to visualize scRNA data.

### Genotyping and genotype data processing

Genotyping was performed using the Infinium Asian Screening Array (Illumina, San Diego, CA), with genotypes called using gtc2vcf plugin on bcftools version 1.19 [33]. PLINK 2 was utilized for genotype preprocessing [34]. Before imputation, we excluded samples with a call rate <0.97, unmatched sex information, or KING-robust kinship estimates <0.08. Variants were excluded if call rate <0.95, minor-allele frequency (MAF) <0.05, or Hardy-Weinberg equilibrium *P* < 1 × 10^-5^. Genotype imputation was performed using the TOPMed Imputation Server with the v2 panel [21]. After imputation, credible variants with imputation quality score *R^2^* > 0.3 were acquired and the same criteria used for pre-processing quality control were re-applied, resulting in 7,779,214 variants.

### Data preprocessing and *cis-*eQTL mapping

We preprocessed genes in the eight major PBMC clusters using a gene count aggregation-based approach (*i.e.*, pseudobulk), as described previously [35]. Before calculating pseudobulk gene expression, genes that were present in less than 1% of all evaluated cells or in fewer than three cells per cluster and sample were removed.

Filtered gene expression was then aggregated based on average SCTransform-normalized counts [22]. After aggregation, we transformed the aggregated values to a normal distribution with mean of 0 and standard deviation of 1.

*cis-*eQTLs were mapped using tensorQTL within 1 Mb windows centered on the transcription starting site (TSS) [23]. The linear regression model used for QTL mapping included 16 covariates: three principal components (PCs) derived from germline variants using PLINK 2, ten PCs derived from the normalized expression matrices, age, and sex as a categorical variable, and scRNA-seq batch information.

### Evaluating condition-specific sc-eQTLs

To estimate the significance of the identified sc-eQTLs across multiple conditions and compare results with the 1M-scbloodNL sc-eQTL study, the multivariate adaptive shrinkage implemented in *mashr* was applied [14, 24]. Specifically, for intra-study significance estimation, we applied the mash algorithm to the most significant eSNP-eGene pairs per each condition, treating these as strong effects and another 100,000 randomly selected eSNP-eGene pairs as random effects. As the sc-eQTL results were inherently sparse, missing data were imputed with a *β* value of 0 and a standard error of 100 to facilitate the significance estimation, and were removed subsequently. The eSNP-eGene pairs selected for further analysis consisted of those with the lowest local false sign rate (lfsr) value for each eGene across all conditions, with consideration of linkage disequilibrium.

Blood-eGenes were annotated as those with a lfsr < 0.05 in any condition. Cell-type eQTLs were defined as those associated with eGenes evaluated in more than five clusters but significant in only one cluster, regardless of ICI treatment. Given our use of adaptive shrinkage, we determined the PosteriorMean and its standard deviation (SD), as well as nominal statistics from tensorQTL [23].

The 1M-scbloodNL study was selected as the replication set as it employs similar cluster annotation and provides the full statistics [14]. We combined the significant eQTLs in each cluster from our dataset (lfsr < 0.05) with those from the untreated condition of the 1M-scbloodNL study (FDR < 0.05), matching the eQTLs based on six major cluster labels. Subsequently, *mashr* was applied using the same parameters as in our analysis.

ICI-treatment-related eGenes were identified by comparing signals between treatments within each cluster using the following criteria: lfsr < 0.05, |Posterior *β*| > 0.5 in one condition, and |Posterior *β* _condition_| / |Posterior *β* _other condition_| > 2.

To identify eGenes specific to our study, we excluded from the total set of significant eGenes any eGene also reported in the GTEx blood eQTL v8 and OneK1K sc-eQTL datasets [12, 13] as significant according to their respective criteria. The resulting study-specific eGenes were used for downstream analysis.

### Functional annotation of eSNPs

Transcript information was obtained from GENCODE version 43 and the distances from candidate eSNPs to TSSs were assessed for both eGenes and non-eGenes. Functional information was annotated using ENCODE candidate *cis*-Regulatory Elements (cCREs) and the Ensembl Variant Effect Predictor (VEP) with database version 109 [36, 37]. Subsequently, we compared the functional annotations of eSNPs with all SNPs considered in our genotype panel to identify those having potential regulatory roles.

### Colocalization analysis

Approximate Bayes factor analysis implemented in coloc v5 (coloc.abf), was utilized to compare the posterior distributions from our eQTL and GWAS studies, which encompassed lung cancer, blood traits, and autoimmune diseases [38] (Additional file 1: Table S7). To estimate the posterior probability of colocalization, harmonized GWAS summary statistics from the GWAS Catalog (https://www.ebi.ac.uk/gwas/) were retrieved and candidate eQTLs with nominal *P* > 1 × 10^-5^ were selected. Due to our focus on the most significant eSNP-eGene pairs identified in the mashr analysis, we used nominal *P*-values and this threshold. After intersecting the GWAS data and eQTLs based on position and alleles, we applied coloc to 1 Mb windows to find possible genetic colocalization, considering a posterior probability (PP.H4) greater than 0.6 as significant. The colocalization results were visualized using the R package *locuszoomR* [39].

### Gene co-expression network analysis using *hdWGCNA*

Weighted gene co-expression network analysis (WGCNA) was conducted using *hdWGCNA* as previously described [25, 28]. Briefly, raw count matrices were utilized to determine gene co-expression networks for each cluster, with parameters set to *k* = 50 and min_cells = 150. Briefly, raw count matrices were utilized to determine gene co-expression networks for each cluster, excluding the Other cluster due to their mixed identity and DC cluster for sparsity. To estimate prognosis-predictive co-expression modules, module expressions between baseline PR and PD groups were compared using the Wilcoxon rank-sum test. To validate our findings, CD8+ T cell-based WGCNA modules were projected onto the AIDA phase 1 dataset and the GSE200996 dataset of ICI-treated HNSCC patients using the ProjecteModules function in *hdWGCNA* [25].

### Gene regulation network analysis using *SCENIC*

Gene regulatory network analysis was conducted using *pySCENIC*, which is a python implementation of single-cell regulatory network inference and clustering (SCENIC) [26]. Due to a technical issue in constructing the gene regulatory network (GRN) with a large number of cells, we first aggregated gene expression data using metacell information from the hdWGCNA analysis and then constructed an adjacency matrix. Next, regulons were identified based on the matrix and cellular enrichment analysis was conducted to annotate regulon activity in each cell. The hdWGCNA networks and pySCENIC regulons were compared based on linear regression of the CD8+ T cell module eigengene score determined using the GetModule function in *hdWGCNA* and the importance score metric from the pySCENIC adjacency matrix. This comparison identified CD8+ T cell-enriched regulons, which received subsequent focus.

### Gene enrichment analysis using GSEA, ChEA3 and *enrichr*

Gene set enrichment analysis (GSEA) was employed to annotate the biological roles of given gene sets [40]. Gene ranks for GSEA were determined using either kME-based gene scores from WGCNA modules or AUC metrics from DEG analysis, based on curated hallmark pathways from the Molecular Signature Database (MsigDB) [41]. Putative transcription factors linked with given WGCNA modules were predicted using ChEA3 based on gene sets from CD8-brown module [42]. The gene enrichment analysis was evaluated using the Kyoto Encyclopedia of Genes and Genomes (KEGG) and Gene Ontology (GO) databases using *enrichr* [43]. Results were considered significant if the false discovery rate (FDR) was less than 0.05.

### Statistical analysis

To evaluate statistical differences of continuous variable between groups, we applied the Wilcoxon rank-sum test and the Kruskal-Wallis test followed by Dunn’s test, or linear regression. For categorical variables, Fisher’s exact test was employed. For survival analysis, the Cox proportional hazard model was utilized. Statistical approaches specific to *cis*-eQTL analysis are described in the eQTL section. Except in *cis*-eQTL mapping and comparison with other eQTL studies, multiple testing corrections were applied using the FDR method. All statistical analyses were conducted using R version 4.3.2.

## Results

### sc-eQTL analysis of PBMC from ICI-treated lung cancer patients

To investigate the influence of germline variants on gene expression in PBMCs from NSCLC patients treated with ICIs, we conducted sc-eQTL analysis on samples from 73 metastatic NSCLC patients given ICI treatment. Samples were collected at baseline and post-treatment (Fig. 1A, Additional file 1: Table S1). After quality assessment steps, 559,714 cells across eight major blood cell populations and 7,779,214 germline variants were acquired (Fig. 1B-E, Additional file 2: Fig. S1). Given the sparse nature of the scRNA-seq data, gene expression values were aggregated (*i.e*., the pseudobulk approach was applied) for the eight major clusters by treatment condition, which identified 9,650 candidate eGenes for eQTL mapping across 16 conditions (Fig. 1C-E). We also mapped *cis*-eQTLs within a 1 Mb of TSSs, employing tensorQTL with a linear model (Fig. 1B-C).

**Figure 1.**
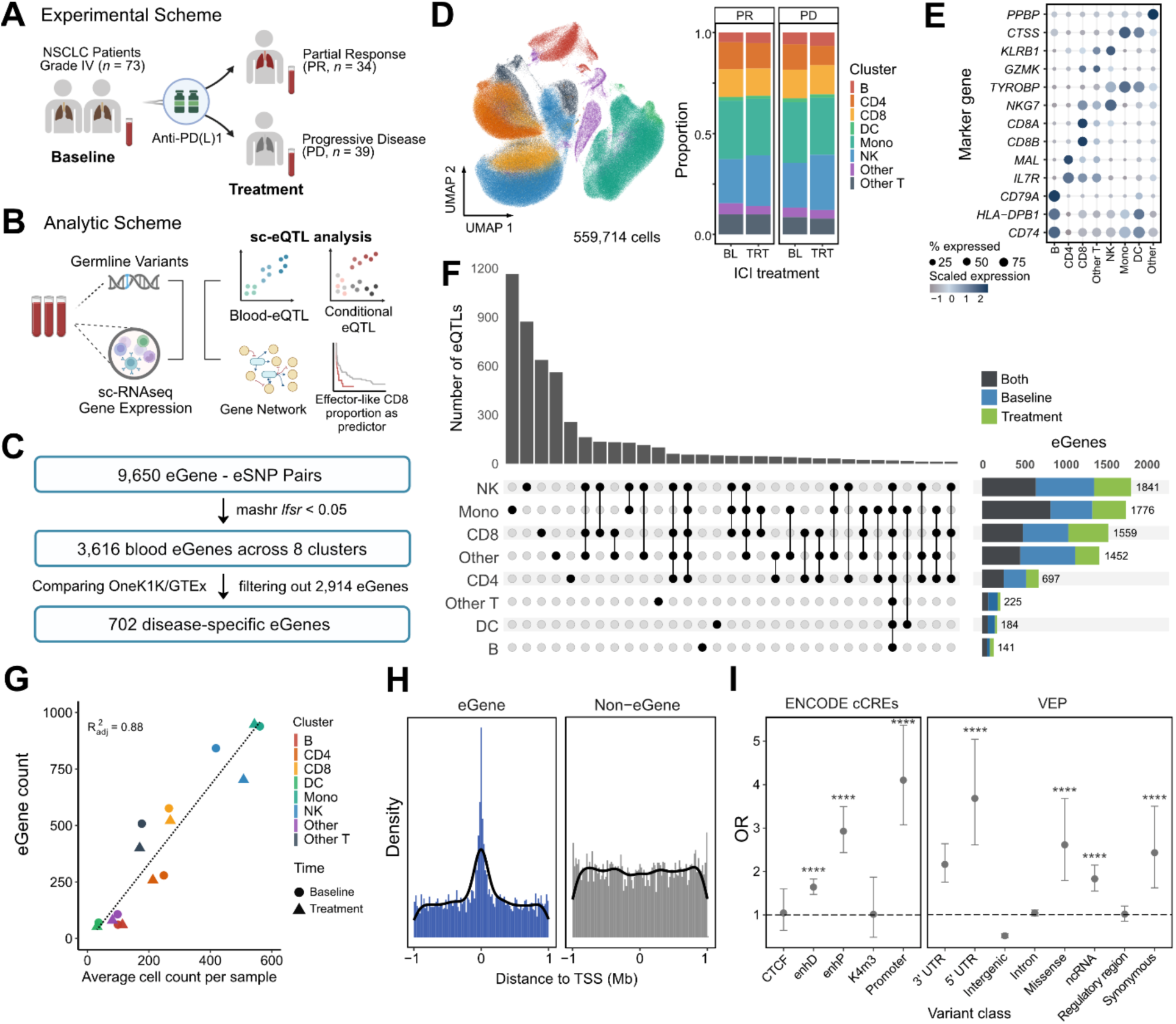
sc-eQTL analysis of circulating immune cells in ICI-treated NSCLC patients. **A** Experimental and **B** analytical schemes. **C** eQTL filtering scheme. **D-E** scRNA-seq analysis of eight major immune cell clusters from PBMC samples of NSCLC patients. **D** UMAP visualizations of immune cell clusters, with cell composition plotted by ICI treatment (BL: baseline, TRT: post-treatment). **E** Dot plot showing cell-type-specific marker expression. **F** UpSet plot of sc-eQTL mapping results, with treatment-based sharing patterns illustrated in the adjacent bar plot. **G** Correlation of average cell count per sample with the number of eGenes identified. Dotted line denotes the regression line. **H** Distribution of distances from TSS for top eGene-eSNP pairs, stratified using lfsr < 0.05. **I** Functional annotation of eSNPs using ENCODE cCREs and Ensemble VEP (****: *P* < 1 x 10^-4^, and error bars represent 95% confidence intervals). enhD and enhP denotes distal and proximal enhancers, respectively.

To ensure robust eQTL mapping, the significance of each eQTL signal and patterns shared among signals were estimated across treatments and clusters using the multivariate adaptive shrinkage (*mashr*) method. This yielded 9,147 eQTLs across 3,616 blood eGenes, determined by meeting a local false sign rate (lfsr) threshold of 0.05 (Fig. 1C, 1F, and Additional file 1: Table S2). Notably, the number of eGenes varied significantly across cell clusters and treatments, and strongly correlated with the number of cells per sample (Fig. 1G; adjusted *R^2^* = 0.88). Comparison of our results with those from GTEx and OneK1K datasets followed by exclusion of overlapped loci revealed 702 lung cancer-specific eGenes, and we defined these as specific eQTLs (Fig. 1C).

As previously demonstrated, eSNPs revealed proximity to TSSs, leading to further enrichments in promoters, 5’ UTRs, and proximal enhancers (as annotated in ENCODE cCRE and Ensembl VEP) (Fig. 1H and 1I) [12, 44, 45]. We also found marginal enrichment in protein-altering variants and non-coding RNAs, suggesting potential functional roles of these variants.

### Cell type and treatment specificity of sc-eQTL

To pinpoint gene regulatory effects of genetic variants in relation to cell type and ICI treatment, we further stratified functional eQTLs across 16 conditions (Fig. 2A, Additional file 1: Table S3). Comparison of eQTL signal sharing within clusters and treatments revealed cell type to have a greater effect than treatment. Specifically, 80– 95% of signals were shared across treatments, while 50–90% were shared within clusters (Fig. 2A). To explore the cell type-specificity of eQTLs in immune-related genes, we extracted significant immune system and immune checkpoint pathway genes with reference to the KEGG database and examined their behaviors. The heatmap shown in Fig. 2B clearly displays a presence of global eQTLs (*e.g., CDC42*, *HLA-C,* etc), cell-type-specific eQTLs (*e.g., PIK3R1, TRADD,* etc), and treatment-specific eQTLs (*e.g., ARPC4, NCK2*, etc). Notably, eQTLs for *CDC42* and *HLA-C* were detected in almost all conditions. *CDC42*, which encodes a Rho GTPase involved in cell migration, cell cycle, and inflammation was downregulated by the alternative allele of rs1534949 across all clusters, with a pronounced effect in CD8+ T cells (posterior *β* = -0.73; Fig. 2C). Meanwhile, *HLA-C*, crucial for self-and viral-antigen recognition with implications in psoriasis via secondary interactions, showed expression differences in CD4+ T cells of 0.73-to 1.25-fold according to genotype at rs9380238 (Fig. 2C) [46].

**Figure 2.**
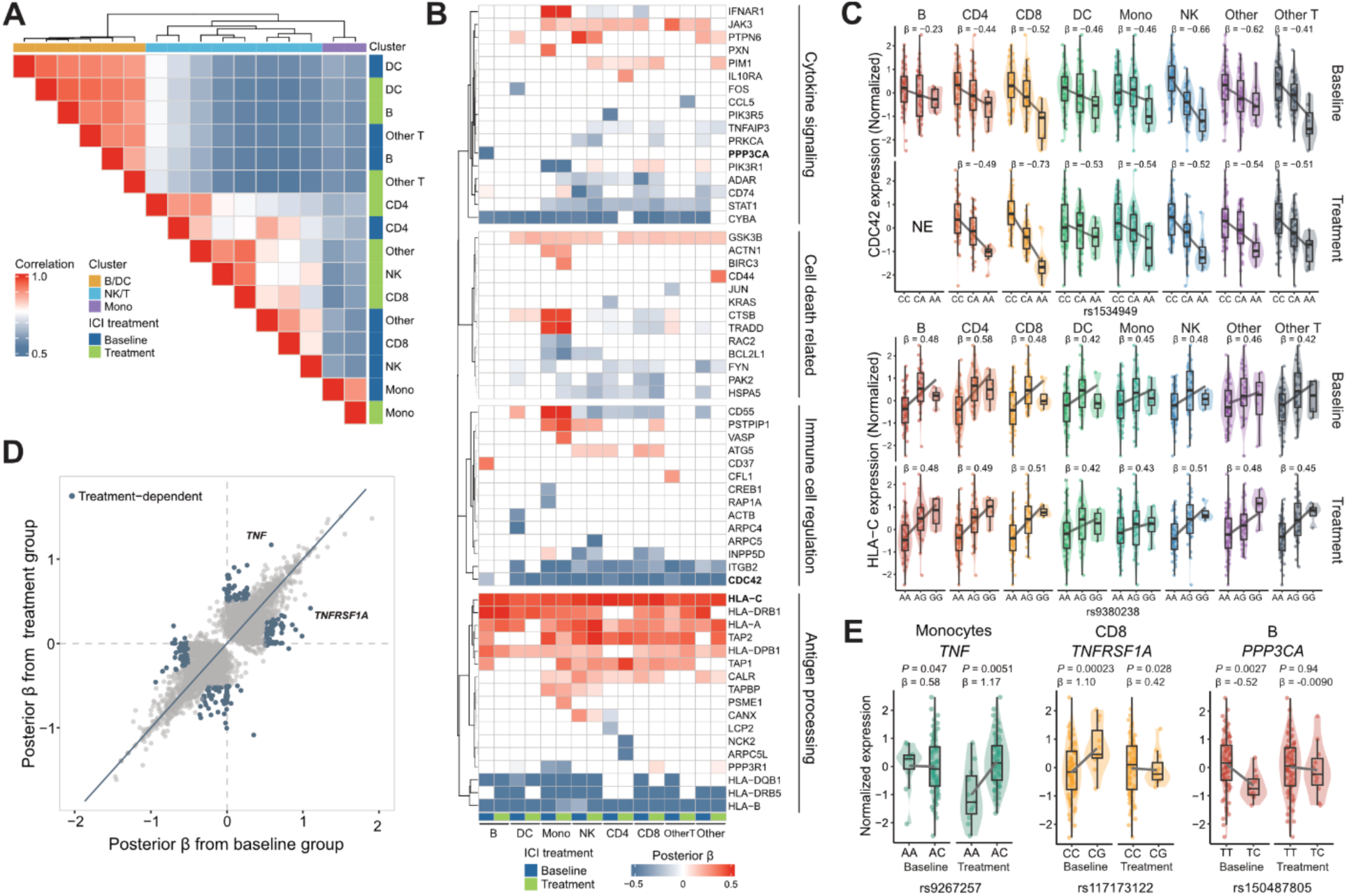
eGenes affect immune cells in cell-type-and treatment-specific manners. **A** Heatmap of sign-sharing rates for significant eQTLs. Color intensity represents the fraction of same-signed signals shared by a pair. **B** Heatmap illustrating treatment-and cluster-based sharing patterns among eGenes from immune system pathways, curated from KEGG. Cell color indicates posterior *β* from mashr analysis. Genes are categorized based on role (right labels). Genes in bold are also displayed in **C** and **E**. **C** Representative eSNP–eGene association plots for eQTLs with effects shared across immune cell clusters and treatment conditions. **D** Scatter plot comparing posterior *β* values from baseline and treatment groups to identify candidate treatment-dependent eQTLs. Blue dots represent the candidate eQTLs. **E** eSNP–eGene association plots displaying ICI treatment-related eQTLs (*i.e.*, response-eQTLs). *P*-values indicate lfsr significance from the mashr analysis. In both (C) and (E), gene expressions is presented as the normalized expression, and posterior *β* values are provided in the plots.

To search for eQTLs whose behavior differs by condition (*i.e*., response-eQTLs), we compared posterior *β* values for each cluster between baseline and treatment; this identified 245 significant ICI treatment-associated eQTLs (Fig. 2D and Additional file 1: Table S4). Examples include *TNF* in monocytes and *TNFRSF1A* in CD8+ T cells (Fig. 2E), which suggest potential genetic modulation of the TNF signaling pathway in cell-cell interactions. Additionally, a *PPP3CA* eQTL in B cells illustrated a cell type-and treatment-specific eQTL effect (Fig. 2E). PPP3CA, also known as calcineurin A, is known for its functions in T cell proliferation [47]. While its functions in B cells are less studied, cyclosporin, an inhibitor of PPP3CA, is reported to directly interfere with the humoral response via B cells [48], suggesting calcineurin regulation to also be important in those cells.

### Validation of eQTL calls using previous eQTL studies

To validate cell-type specificity and context-specific significance of our eQTLs, we compared our findings with existing eQTL datasets from previous studies. First, to validate our sc-eQTL calling strategy and results, we compared our results with the summary statistics from the 1M-scbloodNL study [14] (Fig. 3A; Additional file 1: Table S5). Clusters that corresponded between the two studies shared a larger proportion of eQTLs than did non-corresponding clusters (*P* = 0.002; Fig. 3A, Additional file 2: Fig. S2), supporting the efficacy of our approach.

**Figure 3.**
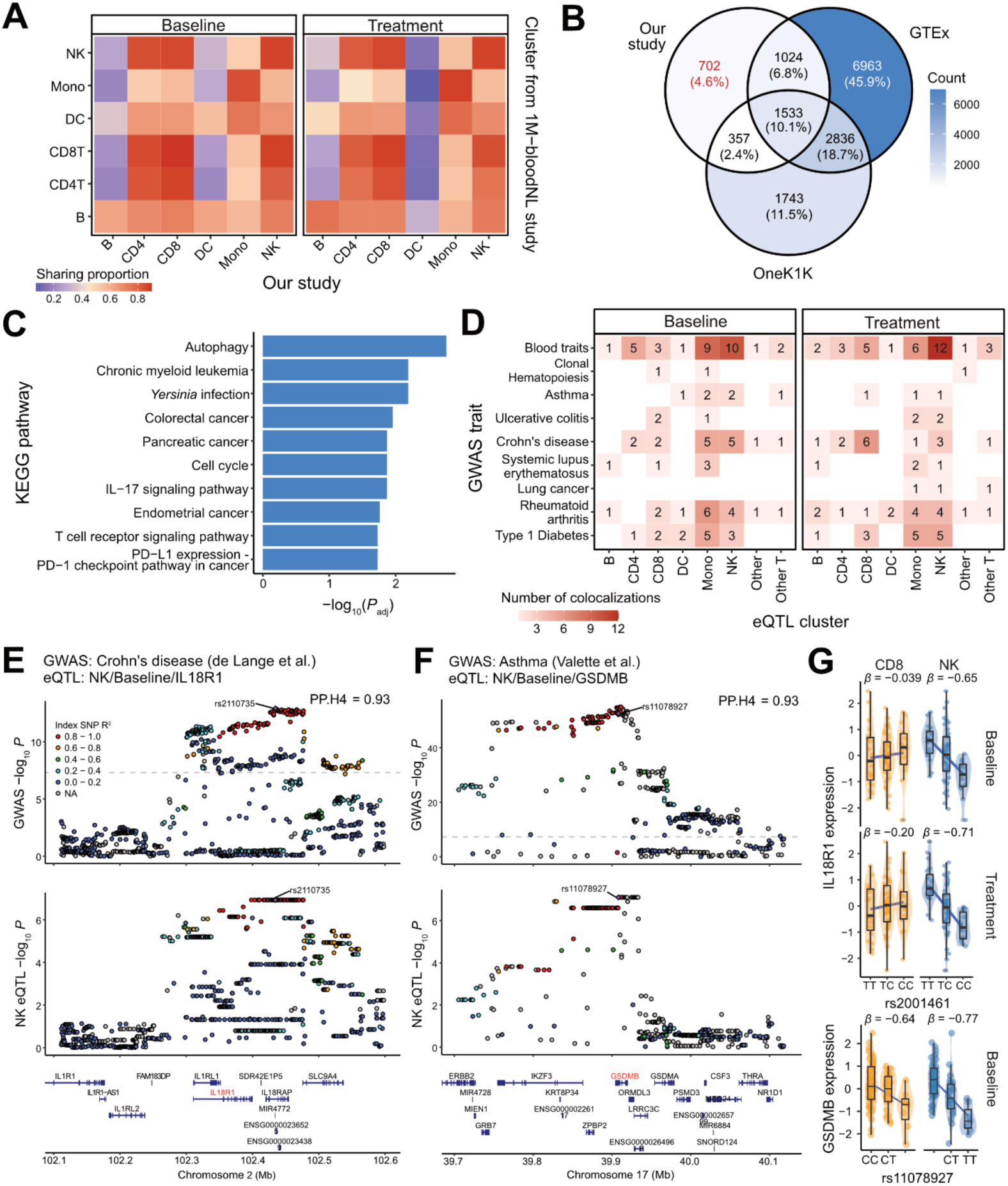
Inter-study eQTL sharing and colocalization with GWAS signals to identify disease-specific pathways and shared autoimmune loci. **A** Heatmap comparing eQTL sharing patterns with the 1M-bloodNL study for both baseline and treatment conditions across major cell clusters, as determined by *mashr* statistics. **B** Venn diagram illustrating the overlap of shared eGenes between our study, GTEx, and the OneK1K study. eQTLs identified exclusively in our study are highlighted in red. **C** KEGG pathway analysis showing enrichment of disease-related pathways in our study-specific eQTLs. The significance level, as adjusted *P*-values were displayed using bar plot. **D** Heatmap displaying the number of colocalizations between eQTL cell clusters and GWAS loci linked to selected disease traits. Colocalizations are shown for both baseline and treatment, with PP.H4 > 0.6 as the threshold. CHIP, clonal hematopoiesis of indeterminate potential; UC, ulcerative colitis; SLE, systemic lupus erythematosus; RA, rheumatoid arthritis; T1D, type 1 diabetes. **E-F** Exemplary eQTL colocalization plots for **E** the *IL18R1* locos in Crohn’s Disease and **F** the *GSDMB* locus in asthma, visualized using *locuszoomR*. The labeled dot on each plot indicates the leading eQTL SNP identified from colocalization analysis. **G** Selected eQTL association plots from **E-F**.

Given the systemic impact of metastatic cancer on the immune system, we hypothesized that the eQTLs identified in our study would exhibit unique eGene profiles, distinct from those observed in other eQTL studies, and hence may illuminate the specific pathophysiological impacts of NSCLC. In this context, we compared our data with representative blood eQTL results from the GTEx consortium and the OneK1K sc-eQTL study in PBMCs. This yielded 702 eGenes unique to our study, which we termed lung cancer-specific eQTLs (Fig. 3B, Additional file 1: Table S6). These eGenes were enriched in disease-related pathways in KEGG, including the PD-L1 pathway and T cell signaling pathways, suggesting germline variants in the tumor-associated immune response (Fig. 3C).

As case studies of this involvement, we present two examples of lung cancer-specific eGenes, *IFNG* and *PTPN11* (Additional file 2: Fig. S3). Interferon-γ is a crucial regulator of immune stimulatory activities in most lineages, but direct control of *IFNG* by germline variants was not reported in previous studies. Given that interferon-γ production by peripheral lymphocytes can determine immune checkpoint responses, the associated variant (rs4913333) may influence patients response to ICI treatment. *PTPN11*, encoding SHP-2, regulates cellular responses to cytokines via the RAS-MAPK pathway; moreover, SHP-2 directly interacts with PD-1 to suppress the immune response of T cells [49]. Interestingly, our eQTL variant rs4767014 near *PTPN11* caused gene expression modulations specific to CD8+ T cells and baseline condition, which may reflect genetic regulation of T cell responses via the PD-1 pathway (Additional file 2: Fig. S3).

### Our eQTL set captures autoimmune trait-related loci in colocalization analysis

Next, we explored the broader genetic context of our condition-and study-specific eQTLs using colocalization analysis on reported GWAS loci related to lung cancer incidence and response to ICI (*e.g.*, blood traits, lung cancer, and autoimmune diseases) (Fig. 3D-G, Additional file 1: Table S7). Our analysis identified 185 loci with significant colocalization probabilities (PP.H4 > 0.6; Fig. 3D, Additional file 1: Table S8). More than 40% of these colocalizations were related to lymphocyte counts and most of the remaining observations were associated with autoimmune diseases, emphasizing the contributions of immune cells in our samples. However, colocalization with lung cancer GWAS signals was less pronounced, possibly because lung cancer pathogenesis and ICI response have distinct genetic etiologies (Fig. 3D).

Notable examples of colocalizations include the NK cell-specific *IL18R1* eQTL with Chron’s disease GWAS locus (Fig. 3E). IL18 is crucial for NK cell activation and proliferation in lung cancer patients, suggesting that the SNP may regulate NK cell activity through IL18 signaling (PP.H4 = 0.93; Fig. 3E and 3G). Additionally, we elucidated a pleiotropic association involving *GSDMB*, which is implicated in multiple autoimmune conditions including asthma, rheumatoid arthritis, and type 1 diabetes (PP.H4 = 0.93 for asthma; Fig. 3F-G, Additional file 2: Fig. S4A). The role of GSDMB in cytolysin highlights its involvement in NK cell cytotoxicity [50]. Moreover, rs4930155 is a *BAD* eQTL specific to treated CD8+ T cells and strongly linked with rs1199047, which shows a strong colocalizations with Crohn’s disease GWAS locus [51] (GWAS *P* = 3.17 x 10^-7^, PP.H4 = 0.79; Additional file 2: Fig. S4B). *BAD* encodes a pro-apoptotic Bcl-2 protein and was not reported as having an associated eQTL in previous studies. Given the role of this gene in apoptosis and immunotherapy, this observation indicates functional regulation of CD8+ T cells by germline variants.

### Identification of CD8+ T effector signature as a predictor of ICI prognosis via gene network analysis

While our eQTL analysis identifies specific genetic variants that regulate the genes related in immune responses and disease-related pathway, understanding their regulatory potential requires examining how these variants influence gene networks within specific cell types or disease contexts. Therefore, we next investigated the co-expression networks to elucidate the broader roles of the eQTLs on immune regulation and ICI prognosis in NSCLC patients. Using *hdWGCNA* and enrichment analyses, we identified ICI-response-predictive networks and their core genes within our dataset (Fig. 4A, Additional file 1: Table S9-S10). The brown module in CD8+ T cells (henceforth referred as “CD8-brown”) exerted the largest effect on module expression differences between responder and non-responder groups at the baseline (Fig. 4A). Gene set analysis revealed this module to be associated with effector functions and antigen-mediated pathways critical for ICI response (Fig. 4B). Module expression is decreased in better prognosis group (PR) (Fig. 4C). Central eGenes in this module, such as *PRF1*, *APOBEC3G*, and *GZMB*, were also downregulated in the group with better prognosis (Additional file 2: Fig. S5).

**Figure 4.**
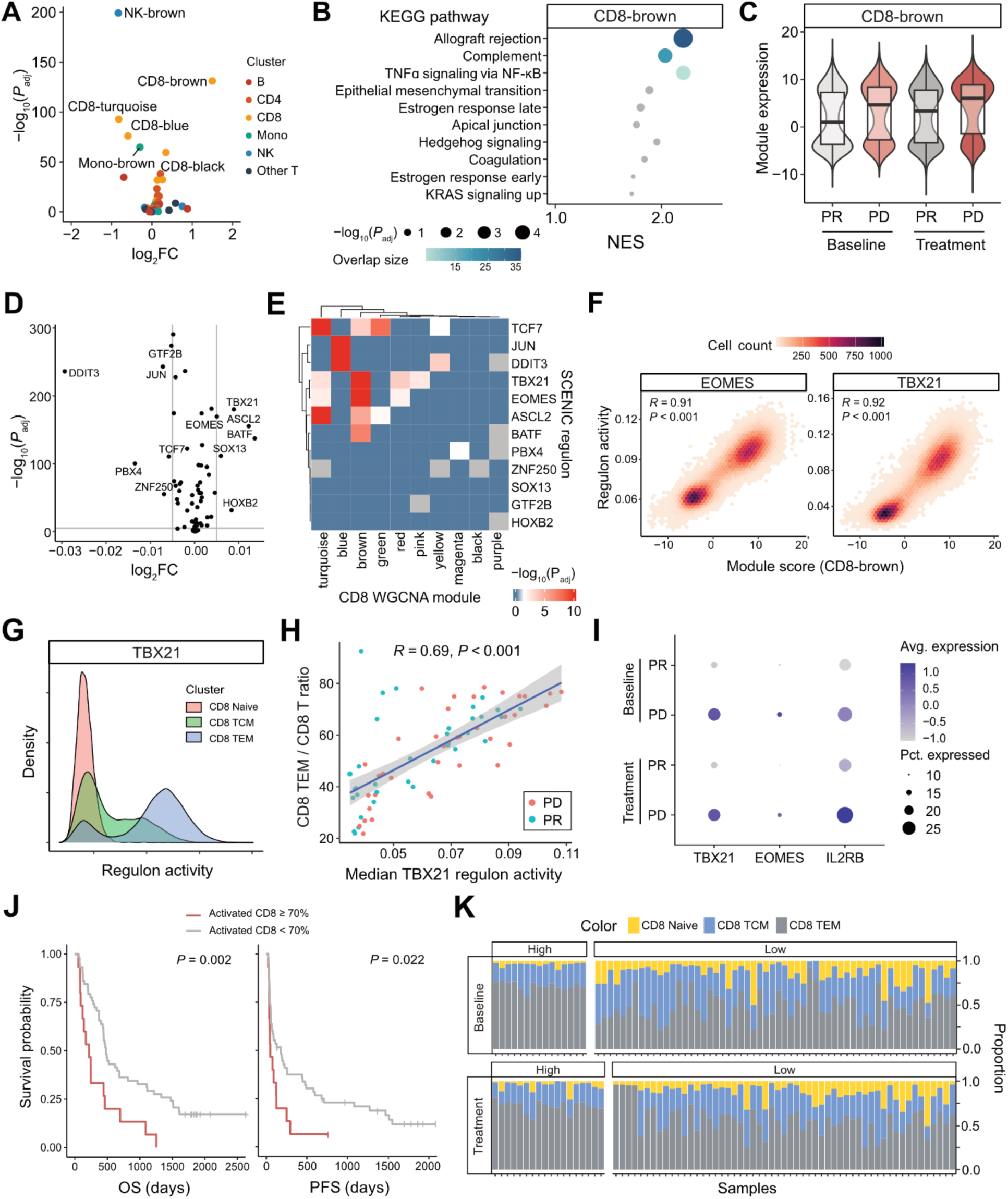
Analysis of disease-responsive gene networks identified in ICI-treated NSCLC patients in relation to CD8+ activation. **A** Volcano plot showing differential module expression between ICI response groups. Colors represent the cell type clusters used to build WGCNA networks. **B** Dot plot contextualized the top GSEA results in relation to msigDB hallmark pathways. Colored dots indicate significant pathway overlap, with dot size representing significance. **C** Violin plot shows expression of the module according to prognosis group and treatment status. **D** Volcano plot illustrating differential CD8-associated gene regulatory networks identified by SCENIC. Labeled regulons indicate those with differential activity between ICI response groups at baseline. **E** Heatmap depicting the correlation of kME from WGCNA with the importance metric from the SCENIC adjacency matrix. Colors indicate the adjusted correlation *P*-value from linear regression. **F** Correlation analysis of the CD8-brown module expression with the TBX21 and EOMES regulons in CD8+ T cells. **G** Density plot showing activity of the TBX21 regulon across CD8+ T cell subsets. **H** Scatter plot showing the correlation of median TBX21 activity with the ratio of CD8+ TEM to total CD8+ T cells in our cohort. **I** Dot plot displaying expression of selected genes across ICI treatment and response groups. **J** Progression-free survival (PFS) and overall survival (OS) plots based on the ratio of activated CD8+ T to total CD8+ T cells, with activation defined based on expression of the CD8-brown module. **K** Bar plots illustrating the breakdown of CD8+ cells in each sample, stratified by high and low proportion of activated CD8 groups defined in **J**.

To validate the robustness of our co-expression networks, we projected the CD8+ T cell co-expression networks onto other PBMC scRNA-seq datasets. CD8-brown module expression was decreased in the AIDA Phase I dataset of healthy donors than both triple-negative breast cancer and head and neck squamous cell carcinoma (HNSCC) datasets [52–54] (Additional file 2: Fig. S6A-B). Furthermore, in blood samples from HNSCC patients undergoing ICI treatment, the module was significantly activated in patients with medium or lower pathological responses to ICI [52] (*P* = 5.9×10^-9^; Additional file 2: Fig. S6C). These findings suggest that the systemic activation of the effector signature of the CD8+ T transcriptome, as represented by CD8-brown module, may be associated with pathological ICI responses, highlighting its potential as a predictive biomarker for ICI treatment outcomes.

### Prioritizing transcriptional factor based on WGCNA and gene regulatory network using SCENIC

To elucidate putative biological drivers for the co-expression network, we utilized SCENIC to estimate transcription factor (TF)-mediated regulon activity. Factors having CD8+ T cell-specific activities were identified, including stress-related TFs such as DDIT3 and JUN, and T cell activation-related TFs such as TBX21, ASCL2, and EOMES (Fig. 4D). A comparison of the adjacency matrix from SCENIC and CD8+ T cells module scores from WGCNA revealed module-specific enrichment of selected regulons (Fig. 4E), with the TBX21 and EOMES regulons particularly showing significant associations with the CD8-brown module (*P* < 0.001 for both regulons; Fig. 4E-F). These findings were further validated using ChEA3, which identified TBX21 as a top-association of the CD8-brown module gene set (Additional file 1: Table S11) [42].

Given that TBX21 and EOMES are essential for CD8+ T cell differentiation and maintenance, we evaluated their activity across CD8+ T cell sub-clusters. Regulon scores showed variable patterns by cell subtypes, with the highest scores obtained for CD8+ T effector memory (CD8+ TEM) cells (Fig.4G). The mean regulon activity of each sample was significantly associated with the fraction of CD8+ TEM cells (Fig. 4H, *R^2^* = 0.69 and *P* < 0.001). Furthermore, we observed higher enrichment of CD8+ TEM in external cancer datasets (Additional file 2: Fig. S6D). Lastly, expression of *TBX21*, *EOMES,* and *IL2RB* (a downstream gene of EOMES) in CD8+ T cells was elevated in patients unresponsive to ICI (Fig. 4I), emphasizing the potential of the effector signature in CD8+ T cells as a possible biomarker of poor prognosis.

### Prediction of ICI response based on the identified CD8+ T cell effector signature

Given that our network analysis identified the CD8+ T cell effector signature network as a potential biomarker for ICI response in lung cancer patients, we subsequently assessed whether this network could also predict survival outcomes under ICI treatment. This analysis indicated that patients with a high ratio (>70%) of activated CD8+ T cells (CD8-brown module score >1.4) to total CD8+ T cells had poorer prognosis in terms of both progression-free survival (PFS, *P* = 0.022) and overall survival (OS, *P* = 0.002; Fig. 4J-K). Given that the CD8+ T effector signature was characterized by significant enrichment of cytotoxic genes, including the core genes *PRF1* and *GZMB*, the baseline signature status of CD8+ T cells in NSCLC patients may predict prognosis under ICI treatment. However, using CD8+ TEM proportion alone as a predictor did not yield significant predictive power, underscoring the specificity of our transcriptomic-network-based model (Additional file 2: Fig. S7).

### Enrichment of eGenes in core genes of CD8+ T co-expression and regulatory networks

Finally, to gain a deeper understanding of the genetic factors that influence the ICI-predictive systemic immune network to ICI responses, we investigated the role of eQTLs within our network analysis (Fig. 5).

**Figure 5.**
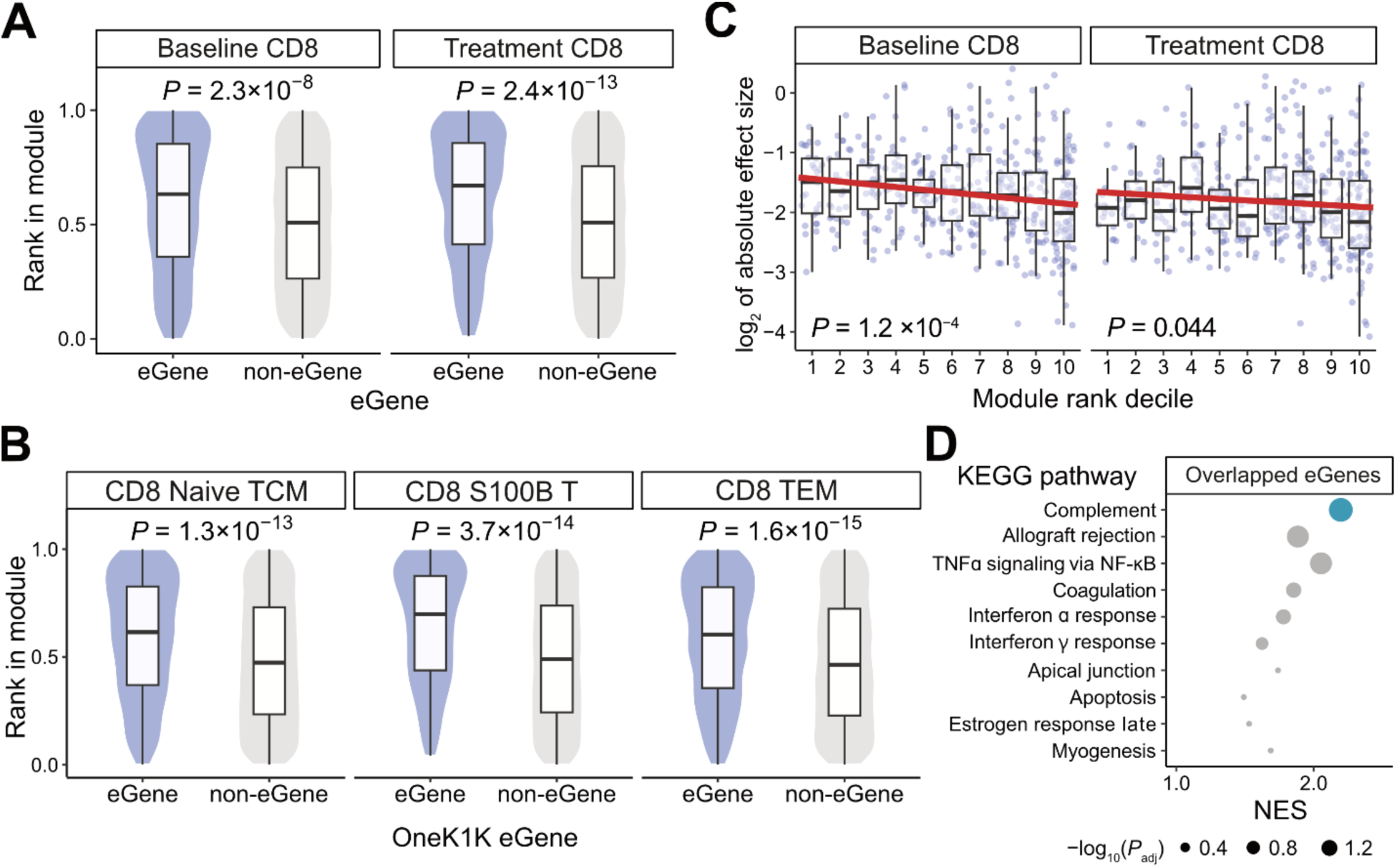
Contribution of eQTLs in response-predictive gene co-expression networks. A-B. Violin plots comparing the rank of eigenvector centrality in CD8+ T cell clusters according to eGene status, **A** in our results and **B** in the OneK1K study. *P*-values were calculated using Wilcoxon’s test. **C** Negative relationship of module centrality by decile rank with eGene effect size in both at baseline and treatment conditions. **D** GSEA results for eGene within the CD8-brown module, showing significant enrichment in the complement pathway. Dot size indicates adjusted *P*-value and dot color significant enrichment.

Using eigenvector-based centrality as a measurement of gene importance within CD8+ T cell WGCNA networks, we found eGenes from our study to exhibit higher centrality compared to non-eGenes (Fig. 5A). This was further supported by SCENIC regulon analysis in CD8+ T cells, which likewise indicated eGenes to be of elevated importance (Additional file 2: Fig. S8). Additionally, eGenes identified in the OneK1K study also displayed higher centrality, emphasizing the role of these shared eGenes as hub genes in our network analysis (Fig. 5B).

Furthermore, we observed negative correlations between eQTL effect size and module ranks both in baseline and treatment conditions (Fig. 5C). Given that core genes often play critical roles within their networks, this suggests that while germline variants broadly influence the immune system through gene expression, their effects on core genes are more constrained, likely due to the tight regulation of network participants. Consistent with this, GSEA analysis of eGenes within the CD8-brown module revealed an enrichment of the complement pathway, which was a shared and centric pathway in the network (Fig. 4B and 5D).

## Discussion

We applied sc-eQTL mapping and transcriptomic network analysis to elucidate the genetic mechanisms underlying regulation of systemic immunity in NSCLC patients given ICI treatment. With this approach, we identified the following: (i) cell type-and treatment-specific gene regulation, (ii) prognosis-predicting gene networks involving TBX21 and EOMES, and (iii) the implications of eQTLs on modulating the strength of gene networks (*e.g.*, CD8-brown). More specifically, our results suggest that PBMC-derived sc-eQTLs in NSCLC patients exhibit distinct cell-type-and treatment-specific signatures. Furthermore, these findings linked changes in ICI-prognosis predicting network to eQTL, thereby implying that the contributions of germline variants to CD8+ T effector signatures may allow for prediction of patient prognosis under ICI treatment.

Further proving the network to identify germline variants with strong effects was not successful, supporting the notion that response to ICI is a complex polygenic trait. Previous eQTL studies in lung cancer have primarily utilized tumor samples or unaffected lung tissues to fine-map GWAS signals or germline mutations, providing functional annotations for lung cancer risk loci [16–18]. Notably, a tumor microenvironment eQTL study using public datasets identified eQTLs in the context of multiple cancers and suggested potential application of eQTL data in predicting risks of melanoma and prostate cancer [55].

As scRNA-seq enables the dissection of immune system heterogeneity, recent eQTL studies on immune cells have leveraged single-cell transcriptome data to elucidate alteration of gene regulation in response to viral load, external perturbations, and disease status [14, 56, 57]. Although each study used a different eQTL mapping approach, both prior and our results consistently identified cell-type-specific and treatment-specific gene regulation, largely dependent on cell-type abundancy (Fig. 1G, 2A). Furthermore, when comparing eGenes with external datasets from healthy donors, our study-specific eGenes (*i.e.* lung cancer eGenes) were comparatively enriched in genes related to ICIs, including *BAD*, *TP53* and *IL1B*, highlighting the potential influence of germline variants in determining the efficacy of ICI treatment. (Fig. 3C, Additional file 1: Table S6).

We further utilized gene co-expression network analysis to identify groups of related genes that potentially predict patient response to ICIs and subsequent prognosis. Among modules identified from the major immune cell populations, we focused on the CD8-brown module due to its potential involvement in ICI treatment (Fig. 4A-D, Additional file 2: Fig. S6C). Regulon analysis linked this module with the TFs EOMES and TBX21, which represent activity of effector cells (Fig. 4E–G). Finally, we determined that the fraction of the CD8+ T cells with this module successfully predicted survival outcome following ICI treatment (Fig. 4J).

Leveraging projection approaches on a healthy control and cancer datasets, we identified activation of cancer-patient-specific CD8+ effector signatures. Centrality analysis revealed eGenes within a module to have higher centrality, suggesting that regulation of gene expression via sc-eQTL is connected within gene regulatory network (Fig. 5, Additional file 2: Fig. S8).

While our study successfully identified cell type-and response-specific gene regulation using the pseudobulk approach with adaptive shrinkage, the sparsity of scRNA-seq data and relatively modest number of samples potentially limited our statistical power. In particular, our analysis of treatment-specific-eQTLs using a linear mixed model identified a substantially lower number of eQTLs than the linear model we presented, highlighting the need for a larger dataset to model interaction effects (Additional file 1: Table S12; 36 eQTLs were detected across all conditions). Nevertheless, our results add additional evidence that the ICI response is a complex trait that are regulated by many genomic variants with small effects.

## Conclusions

In this study, we identified 3,616 blood-and 702 lung cancer-specific eGenes from sc-eQTL analysis and explored their associations with transcriptomic networks predictive of ICI treatment response. Such cell type-and treatment-dependent gene regulation can serve as a valuable resource for understanding systemic immunity in relation to metastatic NSCLC and ICI treatment. Moreover, that eQTLs are linked to ICI-response-predictive networks suggests genetic variants to operate as broad controllers of gene networks. Future research involving larger numbers of tumor patients may validate our finding that germline variants interact with systemic immunity and help facilitate the development of personalized ICI strategies.

## List of abbreviations

*ICI*: Immune checkpoint inhibitor
*NSCLC*: Non-small cell lung cancer
*TSS*: Transcriptional start site
*PBMC*: Peripheral blood mononuclear cell
*sc-eQTL*: Single-cell expression quantitative trait loci
*GWAS*: Genome-wide association study
*GTEx*: Genotype-Tissue Expression
*PR*: Partial response
*PD*: Progressive disease
*SD*: Stable diseasee
*PFS*: Progression-free survival
*OS*: Overall survival
*cCRE*: Candidate cis-regulatory element
*VEP*: Variant effect predictor
*MSigDB*: Molecular Signatures Database
*GSEA*: Gene set enrichment analysis
*WGCNA*: Weighted correlation network analysis
*SCENIC*: Single-cell regulatory network inference and clustering
*GRN*: Gene regulatory network
HNSCC: Head and neck squamous cell carcinoma

## Acknowledgements

We appreciate the participation of the patients in this study. The authors also acknowledge the Korea Research Environment Open Network (KREONET) service and the usage of the Global Science Experimental Data Hub Center (GSDC) provided by Korea Institute of Science and Technology Information (KISTI).

## Declarations

### Ethics approval and consent to participate

This study investigated human samples and was given the permission of SMC Institutional Review Board (No. 2018-04-048, 2022-01-094). All participants underwent an informed consent process before being enrolled in the study.

### Availability of data and materials

The datasets supporting the conclusions of this article are included within the article and its additional files. The datasets used and/or analyzed during the current study are available from the corresponding author on reasonable request.

### Competing interests

The authors declare that they have no competing interests.

### Funding

This work was supported in part by the grants from the Korean Research Foundation (NRF-2021R1A2C3014067, NRF-RS-2023-00207857 to M. Choi, and NRF-2020R1A2C3006535 to S.-H. Lee), the Korea Health Technology R&D Project through the Korea Health Industry Development Institute (KHIDI) (HR20C0025 to S.-H. Lee), and Future Medicine 20*30 Project of the Samsung Medical Center (SMX1230041 to S.-H. Lee).

### Author contributions

SHL, MC designed the study. HS curated, analyzed and performed validation. HS, WYP, SHL and MC interpreted the results from this study. HS, SHL, and MC wrote the original draft. All authors read and approved the final manuscript.

## Additional Files

Additional File 1:

Additional File 1.xlsx; Supplementary tables S1-12.

**Table S1**. Sample information. **Table S2**. Summary statistics of statistically significant eQTLs. **Table S3**. Pairwise sharing proportion matrix from *mashr* analysis. **Table S4**. Summary statistics from treatment-dependent eQTLs. **Table S5**. Sharing Proportion statistics from comparing between our results and 1M-scBloodNL study. **Table S6**. List of lung cancer-specific eQTLs. **Table S7**. List of GWAS publications used in colocalization analysis. **Table S8**. List of significant colocalizations between our eQTL and GWAS traits. **Table S9**. Gene assignment of WGCNA modules and kME scores. Table S10. Top GO enrichment annotations of WGCNA modules. **Table S11**.

Enrichment test results on CD8 brown module, using ChEA3. **Table S12**. sc-eQTL results from linear mixed model based on sample as random effect.

Additional File 2:

Additional File 2.docx; Supplementary figures S1-S8

## Notes

### Competing Interest Statement

The authors have declared no competing interest.

## References

1. Siegel RL, Giaquinto AN, Jemal A: Cancer statistics, 2024. CA Cancer J Clin 2024, 74:12–49.

2. Sampson JN, Wheeler WA, Yeager M, Panagiotou O, Wang Z, Berndt SI, Lan Q, Abnet CC, Amundadottir LT, Figueroa JD, et al: Analysis of Heritability and Shared Heritability Based on Genome-Wide Association Studies for Thirteen Cancer Types. J Natl Cancer Inst 2015, 107:djv279.

3. Bosse Y, Amos CI: A Decade of GWAS Results in Lung Cancer. Cancer Epidemiol Biomarkers Prev 2018, 27:363–379.

4. Hiam-Galvez KJ, Allen BM, Spitzer MH: Systemic immunity in cancer. Nat Rev Cancer 2021, 21:345–359.

5. Wolf AM, Wolf D, Steurer M, Gastl G, Gunsilius E, Grubeck-Loebenstein B: Increase of Regulatory T Cells in the Peripheral Blood of Cancer Patients1. Clinical Cancer Research 2003, 9:606–612.

6. Almand B, Clark JI, Nikitina E, van Beynen J, English NR, Knight SC, Carbone DP, Gabrilovich DI: Increased production of immature myeloid cells in cancer patients: a mechanism of immunosuppression in cancer. J Immunol 2001, 166:678–689.

7. Charrier M, Mezquita L, Lueza B, Dupraz L, Planchard D, Remon J, Caramella C, Cassard L, Boselli L, Reiners KS, et al: Circulating innate immune markers and outcomes in treatment-naive advanced non-small cell lung cancer patients. Eur J Cancer 2019, 108:88–96.

8. Mazieres J, Drilon A, Lusque A, Mhanna L, Cortot AB, Mezquita L, Thai AA, Mascaux C, Couraud S, Veillon R, et al: Immune checkpoint inhibitors for patients with advanced lung cancer and oncogenic driver alterations: results from the IMMUNOTARGET registry. Ann Oncol 2019, 30:1321–1328.

9. Mathios D, Kim JE, Mangraviti A, Phallen J, Park CK, Jackson CM, Garzon-Muvdi T, Kim E, Theodros D, Polanczyk M, et al: Anti-PD-1 antitumor immunity is enhanced by local and abrogated by systemic chemotherapy in GBM. Sci Transl Med 2016, 8:370ra180.

10. Ruhland MK, Roberts EW, Cai E, Mujal AM, Marchuk K, Beppler C, Nam D, Serwas NK, Binnewies M, Krummel MF: Visualizing Synaptic Transfer of Tumor Antigens among Dendritic Cells. Cancer Cell 2020, 37:786–799 e785.

11. Yoo T, Joo SK, Kim HJ, Kim HY, Sim H, Lee J, Kim H-H, Jung S, Lee Y, Jamialahmadi O, et al: Disease-specific eQTL screening reveals an anti-fibrotic effect of AGXT2 in non-alcoholic fatty liver disease. Journal of Hepatology 2021.

12. Consortium TG: The GTEx Consortium atlas of genetic regulatory effects across human tissues. Science 2020, 369:1318–1330.

13. Yazar S, Alquicira-Hernandez J, Wing K, Senabouth A, Gordon MG, Andersen S, Lu Q, Rowson A, Taylor TRP, Clarke L, et al: Single-cell eQTL mapping identifies cell type-specific genetic control of autoimmune disease. Science 2022, 376:eabf3041.

14. Oelen R, de Vries DH, Brugge H, Gordon MG, Vochteloo M, Ye CJ, Westra H-J, Franke L, van der Wijst MGP, single-cell e Qc, Consortium B: Single-cell RNA-sequencing of peripheral blood mononuclear cells reveals widespread, context-specific gene expression regulation upon pathogenic exposure. Nature Communications 2022, 13:3267.

15. Pintarelli G, Cotroneo CE, Noci S, Dugo M, Galvan A, Delli Carpini S, Citterio L, Manunta P, Incarbone M, Tosi D, et al: Genetic susceptibility variants for lung cancer: replication study and assessment as expression quantitative trait loci. Sci Rep 2017, 7:42185.

16. Nguyen JD, Lamontagne M, Couture C, Conti M, Pare PD, Sin DD, Hogg JC, Nickle D, Postma DS, Timens W, et al: Susceptibility loci for lung cancer are associated with mRNA levels of nearby genes in the lung. Carcinogenesis 2014, 35:2653–2659.

17. McKay JD, Hung RJ, Han Y, Zong X, Carreras-Torres R, Christiani DC, Caporaso NE, Johansson M, Xiao X, Li Y, et al: Large-scale association analysis identifies new lung cancer susceptibility loci and heterogeneity in genetic susceptibility across histological subtypes. Nat Genet 2017, 49:1126–1132.

18. Gong J, Mei S, Liu C, Xiang Y, Ye Y, Zhang Z, Feng J, Liu R, Diao L, Guo AY, et al: PancanQTL: systematic identification of cis-eQTLs and trans-eQTLs in 33 cancer types. Nucleic Acids Res 2018, 46:D971–D976.

19. Bosse Y, Li Z, Xia J, Manem V, Carreras-Torres R, Gabriel A, Gaudreault N, Albanes D, Aldrich MC, Andrew A, et al: Transcriptome-wide association study reveals candidate causal genes for lung cancer. Int J Cancer 2020, 146:1862–1878.

20. Byun J, Han Y, Li Y, Xia J, Long E, Choi J, Xiao X, Zhu M, Zhou W, Sun R, et al: Cross-ancestry genome-wide meta-analysis of 61,047 cases and 947,237 controls identifies new susceptibility loci contributing to lung cancer. Nat Genet 2022, 54:1167–1177.

21. Das S, Forer L, Schonherr S, Sidore C, Locke AE, Kwong A, Vrieze SI, Chew EY, Levy S, McGue M, et al: Next-generation genotype imputation service and methods. Nat Genet 2016, 48:1284–1287.

22. Hafemeister C, Satija R: Normalization and variance stabilization of single-cell RNA-seq data using regularized negative binomial regression. Genome Biol 2019, 20:296.

23. Taylor-Weiner A, Aguet F, Haradhvala NJ, Gosai S, Anand S, Kim J, Ardlie K, Van Allen EM, Getz G: Scaling computational genomics to millions of individuals with GPUs. Genome Biol 2019, 20:228.

24. Urbut SM, Wang G, Carbonetto P, Stephens M: Flexible statistical methods for estimating and testing effects in genomic studies with multiple conditions. Nat Genet 2019, 51:187–195.

25. Morabito S, Reese F, Rahimzadeh N, Miyoshi E, Swarup V: hdWGCNA identifies co-expression networks in high-dimensional transcriptomics data. Cell Rep Methods 2023, 3:100498.

26. Van de Sande B, Flerin C, Davie K, De Waegeneer M, Hulselmans G, Aibar S, Seurinck R, Saelens W, Cannoodt R, Rouchon Q, et al: A scalable SCENIC workflow for single-cell gene regulatory network analysis. Nat Protoc 2020, 15:2247–2276.

27. Kim H, Park S, Han KY, Lee N, Kim H, Jung HA, Sun JM, Ahn JS, Ahn MJ, Lee SH, Park WY: Clonal expansion of resident memory T cells in peripheral blood of patients with non-small cell lung cancer during immune checkpoint inhibitor treatment. J Immunother Cancer 2023, 11.

28. Sim H, Park HJ, Park G-H, Kim YJ, Park W-Y, Lee S-H, Choi M: Increased inflammatory signature in myeloid cells of non-small cell lung cancer patients with high clonal hematopoiesis burden. eLife 2024:RP96951.

29. Kang HM, Subramaniam M, Targ S, Nguyen M, Maliskova L, McCarthy E, Wan E, Wong S, Byrnes L, Lanata CM, et al: Multiplexed droplet single-cell RNA-sequencing using natural genetic variation. Nat Biotechnol 2018, 36:89–94.

30. Fleming SJ, Chaffin MD, Arduini A, Akkad AD, Banks E, Marioni JC, Philippakis AA, Ellinor PT, Babadi M: Unsupervised removal of systematic background noise from droplet-based single-cell experiments using CellBender. Nat Methods 2023, 20:1323–1335.

31. Hao Y, Hao S, Andersen-Nissen E, Mauck WM, 3rd, Zheng S, Butler A, Lee MJ, Wilk AJ, Darby C, Zager M, et al: Integrated analysis of multimodal single-cell data. Cell 2021, 184:3573–3587 e3529.

32. Korsunsky I, Millard N, Fan J, Slowikowski K, Zhang F, Wei K, Baglaenko Y, Brenner M, Loh PR, Raychaudhuri S: Fast, sensitive and accurate integration of single-cell data with Harmony. Nat Methods 2019, 16:1289–1296.

33. Danecek P, Bonfield JK, Liddle J, Marshall J, Ohan V, Pollard MO, Whitwham A, Keane T, McCarthy SA, Davies RM, Li H: Twelve years of SAMtools and BCFtools. Gigascience 2021, 10.

34. Chang CC, Chow CC, Tellier LCAM, Vattikuti S, Purcell SM, Lee JJ: Second-generation PLINK: rising to the challenge of larger and richer datasets. GigaScience 2015, 4:7.

35. Cuomo ASE, Alvari G, Azodi CB, single-cell e Qc, McCarthy DJ, Bonder MJ: Optimizing expression quantitative trait locus mapping workflows for single-cell studies. Genome Biol 2021, 22:188.

36. Consortium EP, Moore JE, Purcaro MJ, Pratt HE, Epstein CB, Shoresh N, Adrian J, Kawli T, Davis CA, Dobin A, et al: Expanded encyclopaedias of DNA elements in the human and mouse genomes. Nature 2020, 583:699–710.

37. McLaren W, Gil L, Hunt SE, Riat HS, Ritchie GR, Thormann A, Flicek P, Cunningham F: The Ensembl Variant Effect Predictor. Genome Biol 2016, 17:122.

38. Wang G, Sarkar A, Carbonetto P, Stephens M: A simple new approach to variable selection in regression, with application to genetic fine mapping. J R Stat Soc Series B Stat Methodol 2020, 82:1273–1300.

39. Pruim RJ, Welch RP, Sanna S, Teslovich TM, Chines PS, Gliedt TP, Boehnke M, Abecasis GR, Willer CJ: LocusZoom: regional visualization of genome-wide association scan results. Bioinformatics 2010, 26:2336–2337.

40. Sergushichev AA: An algorithm for fast preranked gene set enrichment analysis using cumulative statistic calculation. bioRxiv 2016:060012.

41. Liberzon A, Birger C, Thorvaldsdottir H, Ghandi M, Mesirov JP, Tamayo P: The Molecular Signatures Database (MSigDB) hallmark gene set collection. Cell Syst 2015, 1:417–425.

42. Keenan AB, Torre D, Lachmann A, Leong AK, Wojciechowicz ML, Utti V, Jagodnik KM, Kropiwnicki E, Wang Z, Ma’ayan A: ChEA3: transcription factor enrichment analysis by orthogonal omics integration. Nucleic Acids Res 2019, 47:W212–W224.

43. Kuleshov MV, Jones MR, Rouillard AD, Fernandez NF, Duan Q, Wang Z, Koplev S, Jenkins SL, Jagodnik KM, Lachmann A, et al: Enrichr: a comprehensive gene set enrichment analysis web server 2016 update. Nucleic Acids Res 2016, 44:W90–97.

44. Natri HM, Del Azodi CB, Peter L, Taylor CJ, Chugh S, Kendle R, Chung MI, Flaherty DK, Matlock BK, Calvi CL, et al: Cell-type-specific and disease-associated expression quantitative trait loci in the human lung. Nat Genet 2024, 56:595–604.

45. Bryois J, Calini D, Macnair W, Foo L, Urich E, Ortmann W, Iglesias VA, Selvaraj S, Nutma E, Marzin M, et al: Cell-type-specific cis-eQTLs in eight human brain cell types identify novel risk genes for psychiatric and neurological disorders. Nat Neurosci 2022, 25:1104–1112.

46. Lee KY, Leung KS, Tang NLS, Wong MH: Discovering Genetic Factors for psoriasis through exhaustively searching for significant second order SNP-SNP interactions. Sci Rep 2018, 8:15186.

47. Medyouf H, Alcalde H, Berthier C, Guillemin MC, dos Santos NR, Janin A, Decaudin D, de The H, Ghysdael J: Targeting calcineurin activation as a therapeutic strategy for T-cell acute lymphoblastic leukemia. Nat Med 2007, 13:736–741.

48. De Bruyne R, Bogaert D, De Ruyck N, Lambrecht BN, Van Winckel M, Gevaert P, Dullaers M: Calcineurin inhibitors dampen humoral immunity by acting directly on naive B cells. Clin Exp Immunol 2015, 180:542–550.

49. Cai Z, Kotzin JJ, Ramdas B, Chen S, Nelanuthala S, Palam LR, Pandey R, Mali RS, Liu Y, Kelley MR, et al: Inhibition of Inflammatory Signaling in Tet2 Mutant Preleukemic Cells Mitigates Stress-Induced Abnormalities and Clonal Hematopoiesis. Cell Stem Cell 2018, 23:833–849 e835.

50. Hansen JM, de Jong MF, Wu Q, Zhang LS, Heisler DB, Alto LT, Alto NM: Pathogenic ubiquitination of GSDMB inhibits NK cell bactericidal functions. Cell 2021, 184:3178–3191 e3118.

51. de Lange KM, Moutsianas L, Lee JC, Lamb CA, Luo Y, Kennedy NA, Jostins L, Rice DL, Gutierrez-Achury J, Ji SG, et al: Genome-wide association study implicates immune activation of multiple integrin genes in inflammatory bowel disease. Nat Genet 2017, 49:256–261.

52. Luoma AM, Suo S, Wang Y, Gunasti L, Porter CBM, Nabilsi N, Tadros J, Ferretti AP, Liao S, Gurer C, et al: Tissue-resident memory and circulating T cells are early responders to pre-surgical cancer immunotherapy. Cell 2022, 185:2918–2935 e2929.

53. Zhang Y, Chen H, Mo H, Hu X, Gao R, Zhao Y, Liu B, Niu L, Sun X, Yu X, et al: Single-cell analyses reveal key immune cell subsets associated with response to PD-L1 blockade in triple-negative breast cancer. Cancer Cell 2021, 39:1578–1593 e1578.

54. Kock KH, Tan LM, Han KY, Ando Y, Jevapatarakul D, Chatterjee A, Lin QXX, Buyamin EV, Sonthalia R, Rajagopalan D, et al: Single-cell analysis of human diversity in circulating immune cells. bioRxiv 2024.

55. Pagadala M, Sears TJ, Wu VH, Perez-Guijarro E, Kim H, Castro A, Talwar JV, Gonzalez-Colin C, Cao S, Schmiedel BJ, et al: Germline modifiers of the tumor immune microenvironment implicate drivers of cancer risk and immunotherapy response. Nat Commun 2023, 14:2744.

56. Aquino Y, Bisiaux A, Li Z, O’Neill M, Mendoza-Revilla J, Merkling SH, Kerner G, Hasan M, Libri V, Bondet V, et al: Dissecting human population variation in single-cell responses to SARS-CoV-2. Nature 2023, 621:120–128.

57. Perez RK, Gordon MG, Subramaniam M, Kim MC, Hartoularos GC, Targ S, Sun Y, Ogorodnikov A, Bueno R, Lu A, et al: Single-cell RNA-seq reveals cell type-specific molecular and genetic associations to lupus. Science 2022, 376:eabf1970.

