## Supplemental figures for "Systemic CD8+ T cell effector signature predicts prognosis of lung cancer immunotherapy"

**Supplementary Figures**

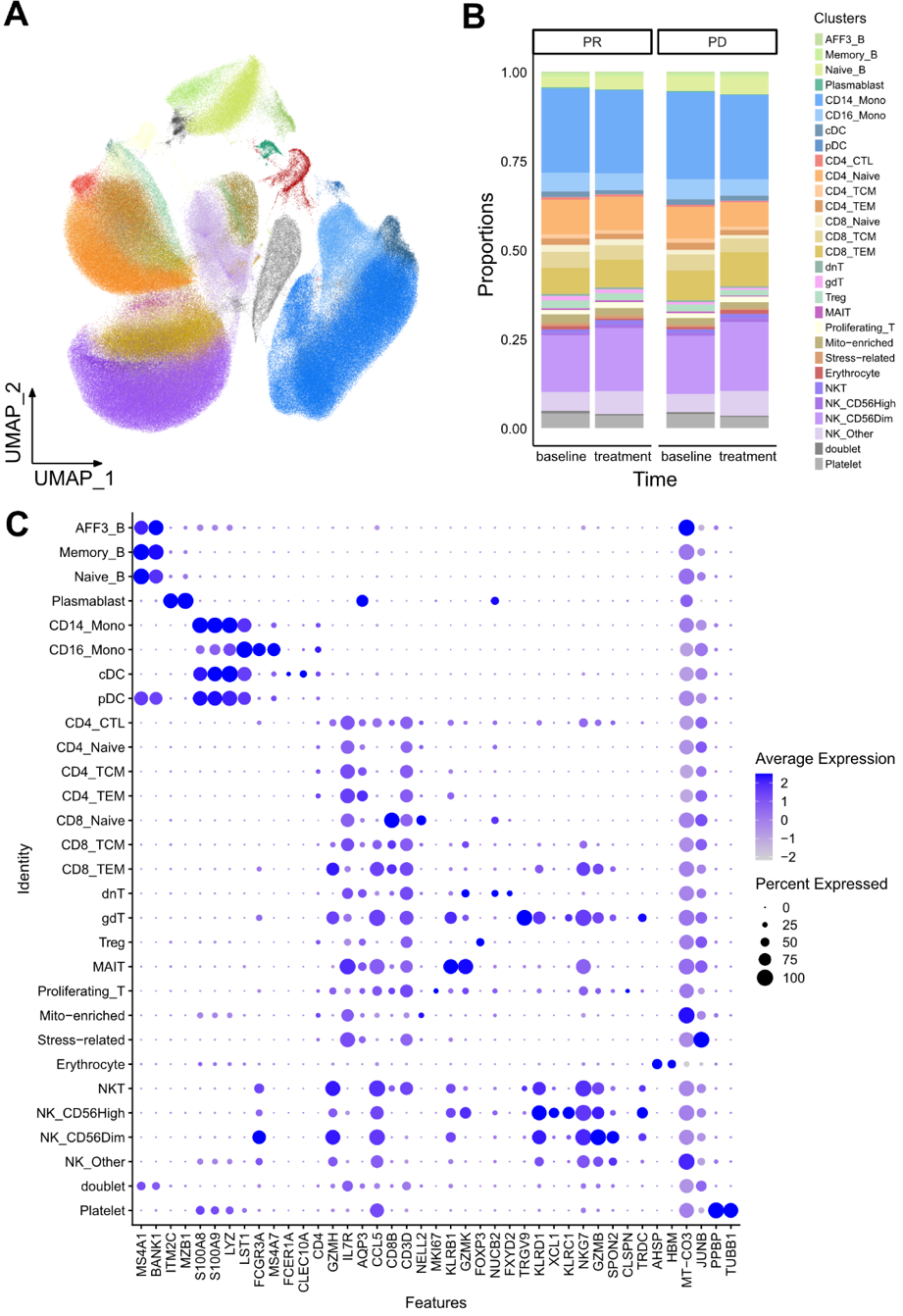

**Supplementary Figure S1. Detailed annotation of PBMC scRNA-seq from ICI-treated NSCLC patients.** **A** Representative UMAP plot using subclusters. **B** Bar Plot using cell proportion by treatment and response groups. **C** Gene Expression markers of subclusters. The color indicates expression levels and size of dots shows percentage of expression in the clusters.

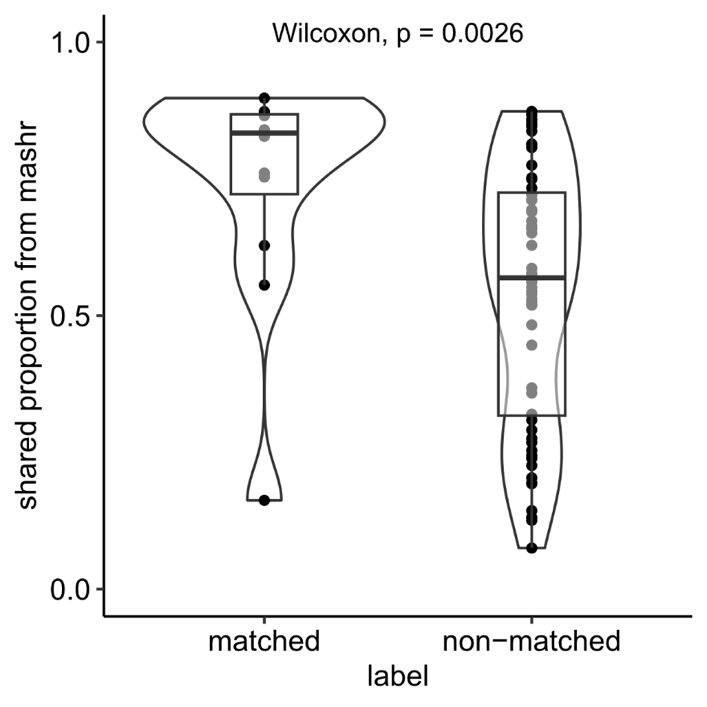

**Supplementary Figure S2.** Distribution of shared proportions between matched and non-matched cluster label from our study and sc-1MbloodNL study.

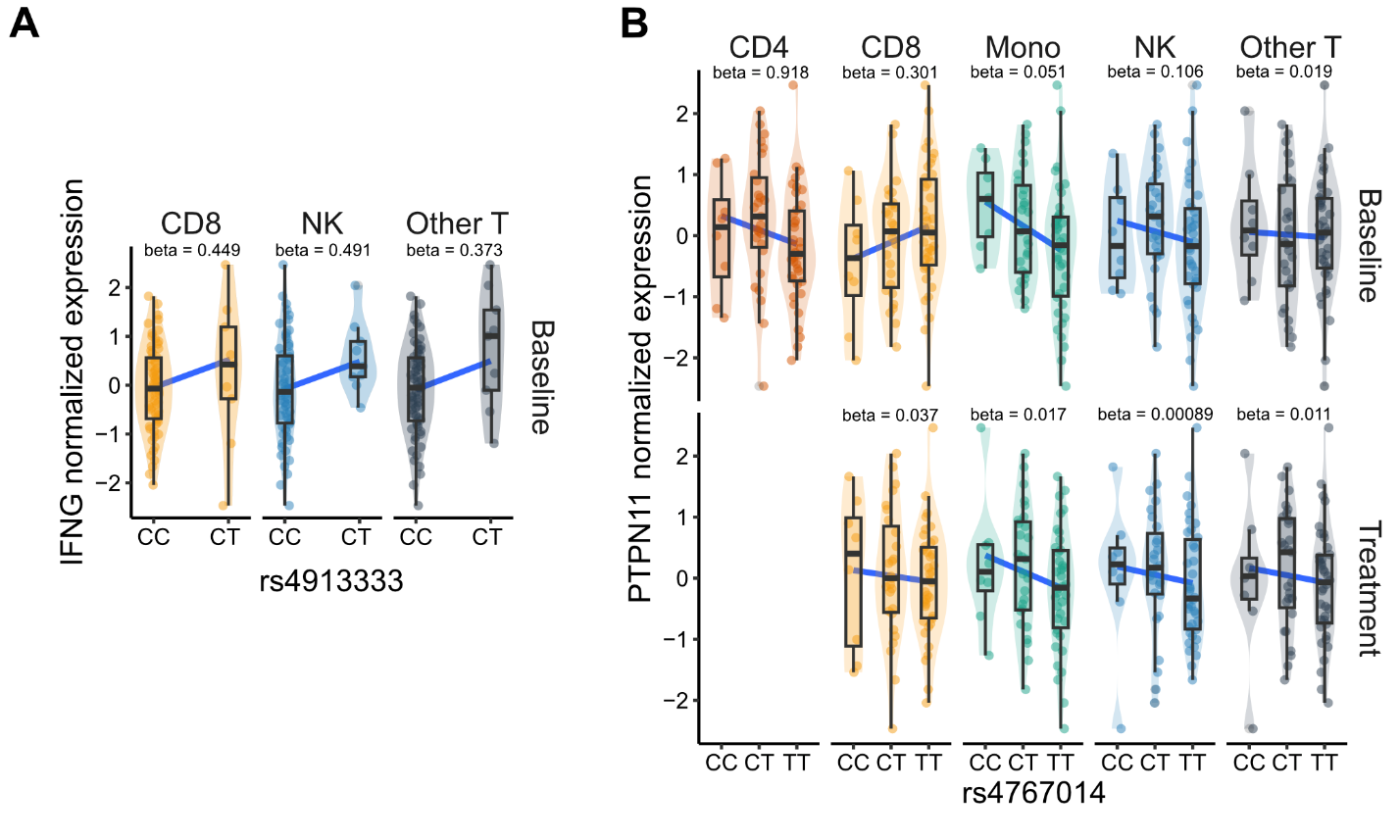

**Supplementary Figure S3.** Examples of Lung cancer specific-eQTL plots in **A** *IFNG* and **B** *PTPN11* genes.

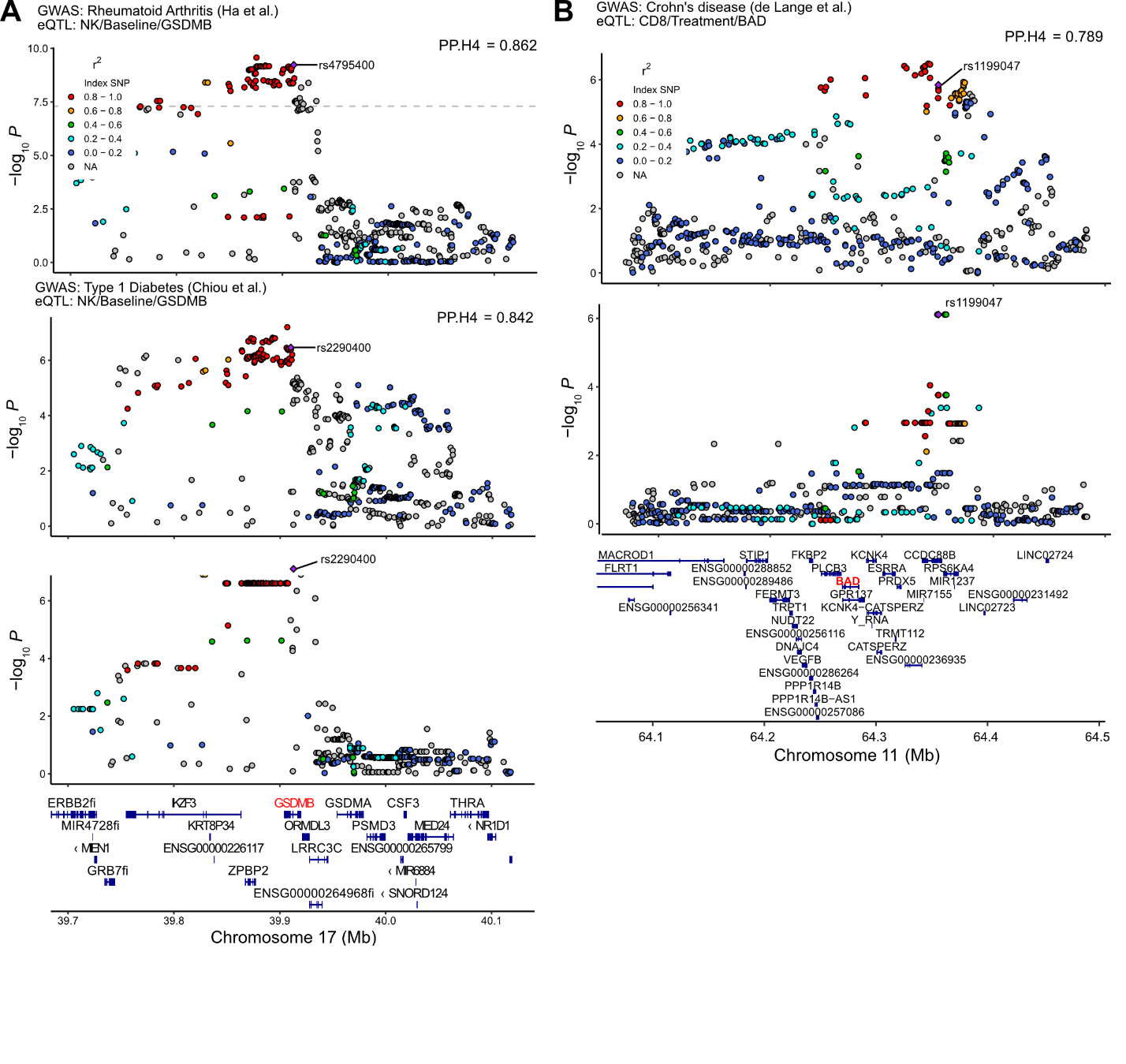

**Supplementary Figure S4.** Colocalization analysis results from **A** pleiotropic associations in auto-immune loci to GSDMB and **B** Crohn’s disease to BAD eQTL.

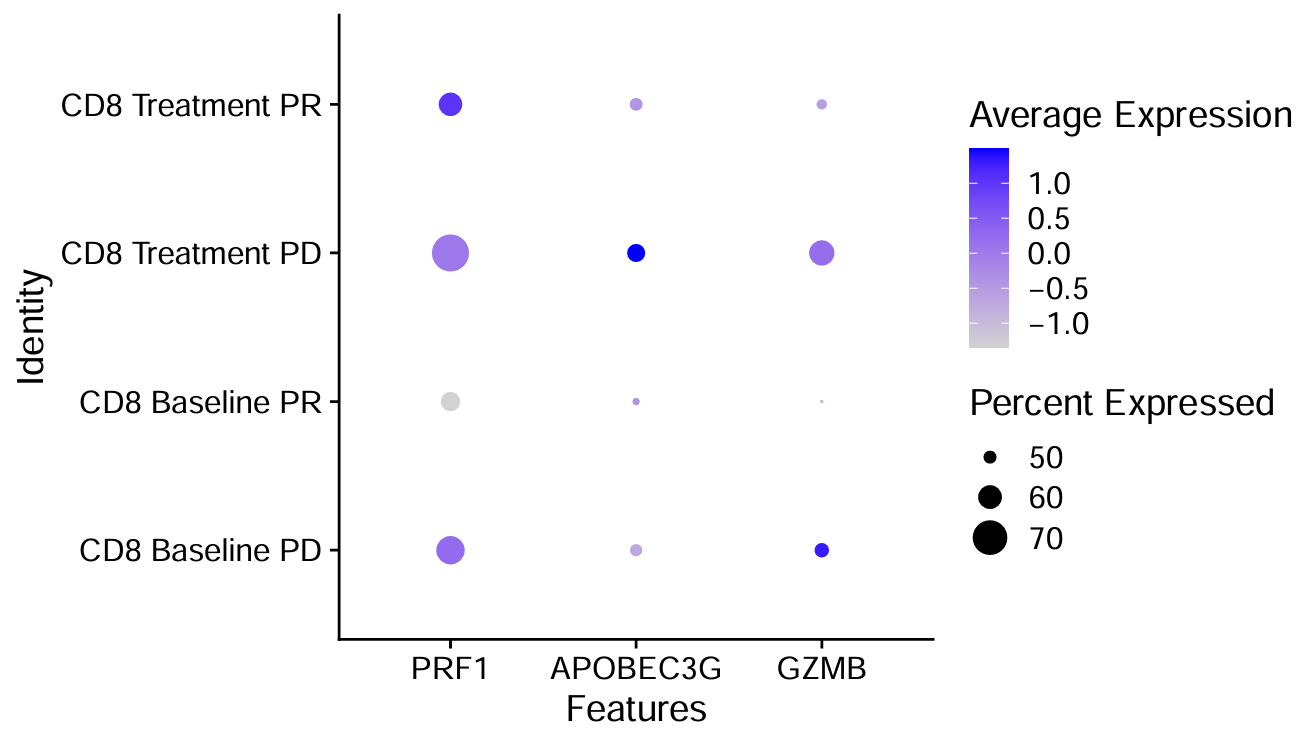

**Supplementary Figure S5.** Core gene expressions in CD8 T Cells in the CD8-brown module, stratified by treatment and response to ICI.

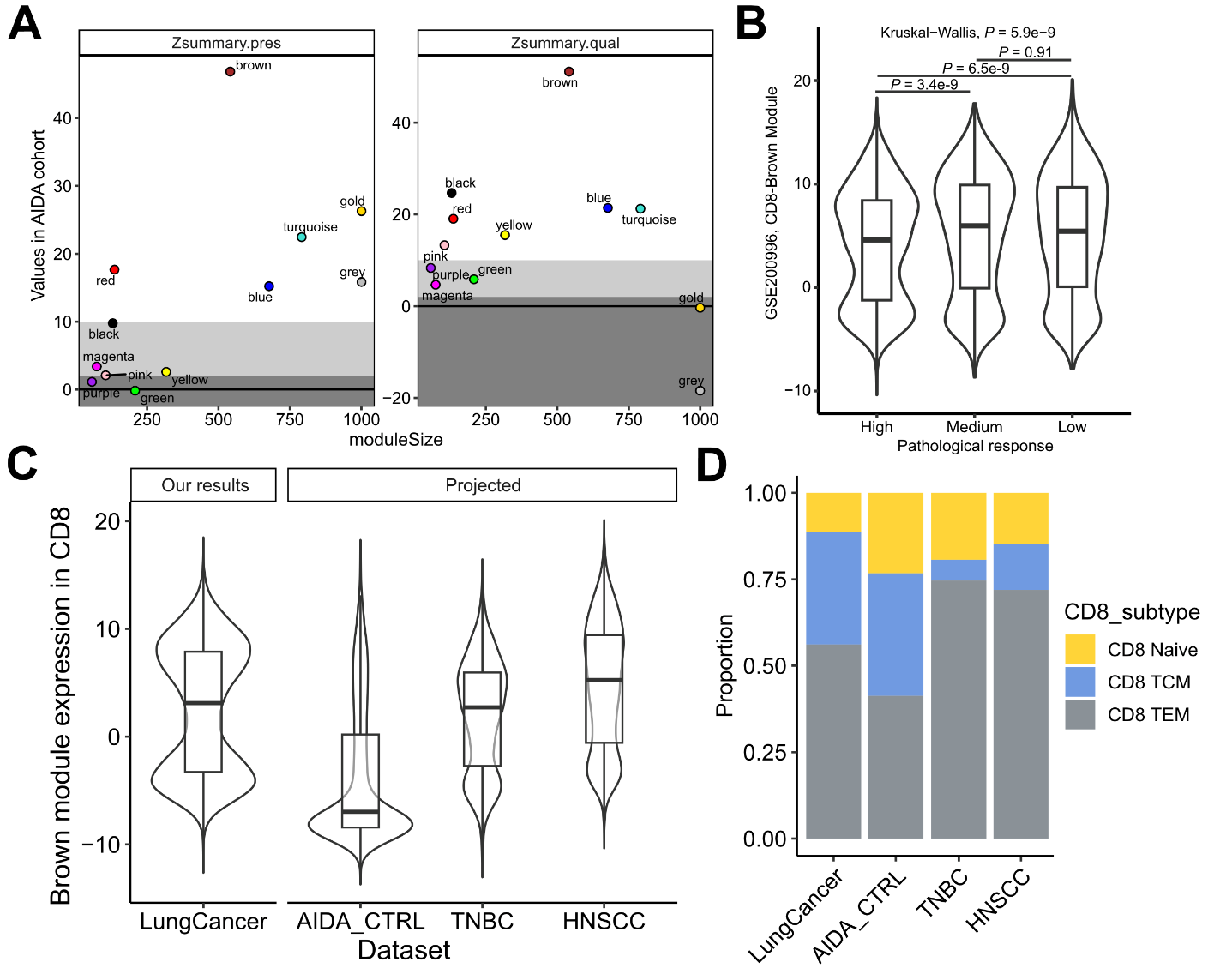

**Supplementary Figure S6. Validation of WGCNA analysis using external datasets.** **A** Projection results of WGCNA modules from CD8 T cells using AIDA dataset. **B** The "brown" module score distribution in external datasets. **C** A density plot based on projected dataset GSE200996 with TNBC patients, by pathological response provided from the study. **D** Estimated CD8 subtypes from external scRNA-seq studies. "LungCancer" indicates our study, with colors indicating CD8 subtypes as defined in Azimuth human PBMC reference.

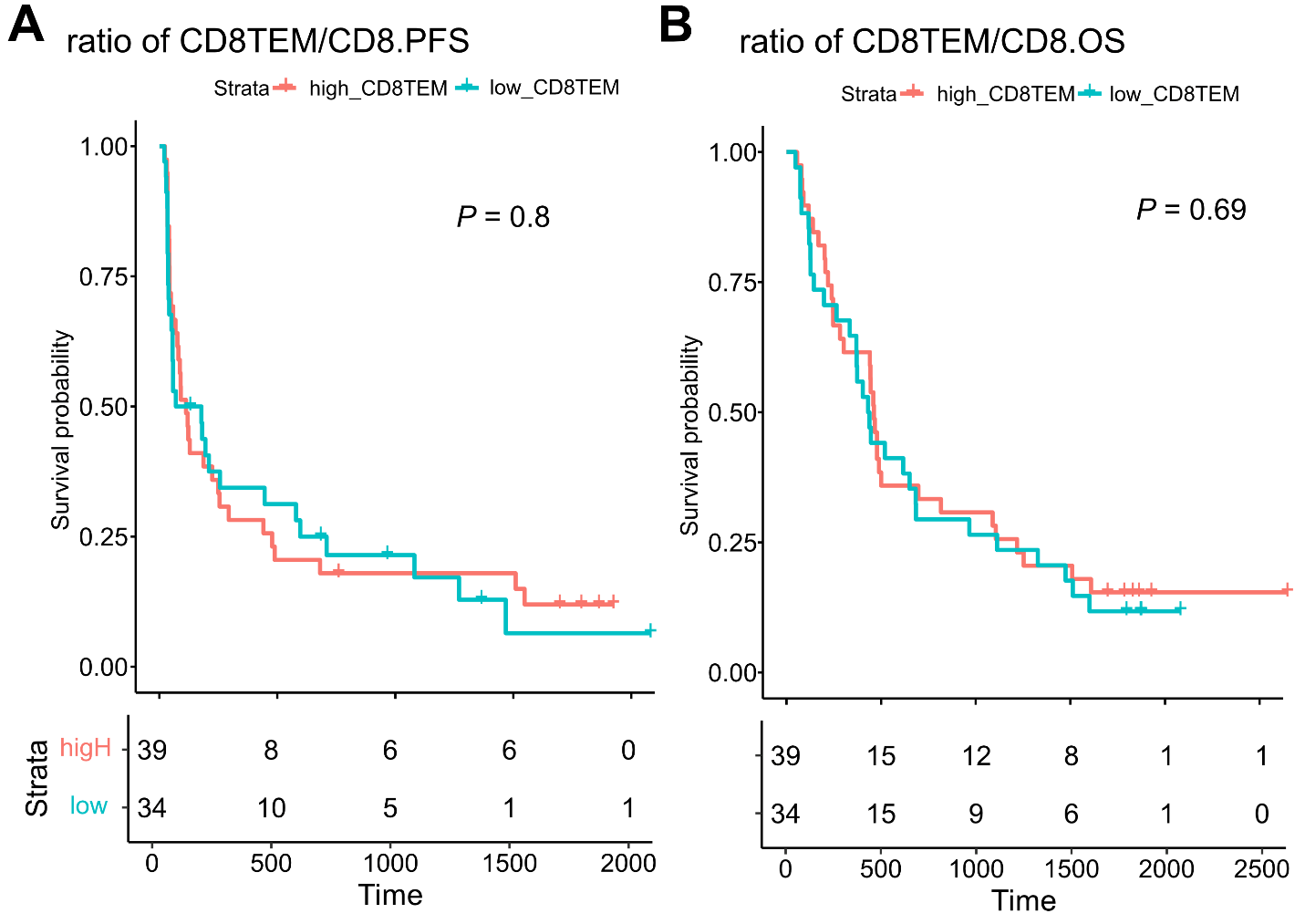

**Supplementary Figure S7.** Survival plot based on CD8 TEM to Total CD8 T cell fractions, showing less predictive powers in **A** PFS and **B** OS.

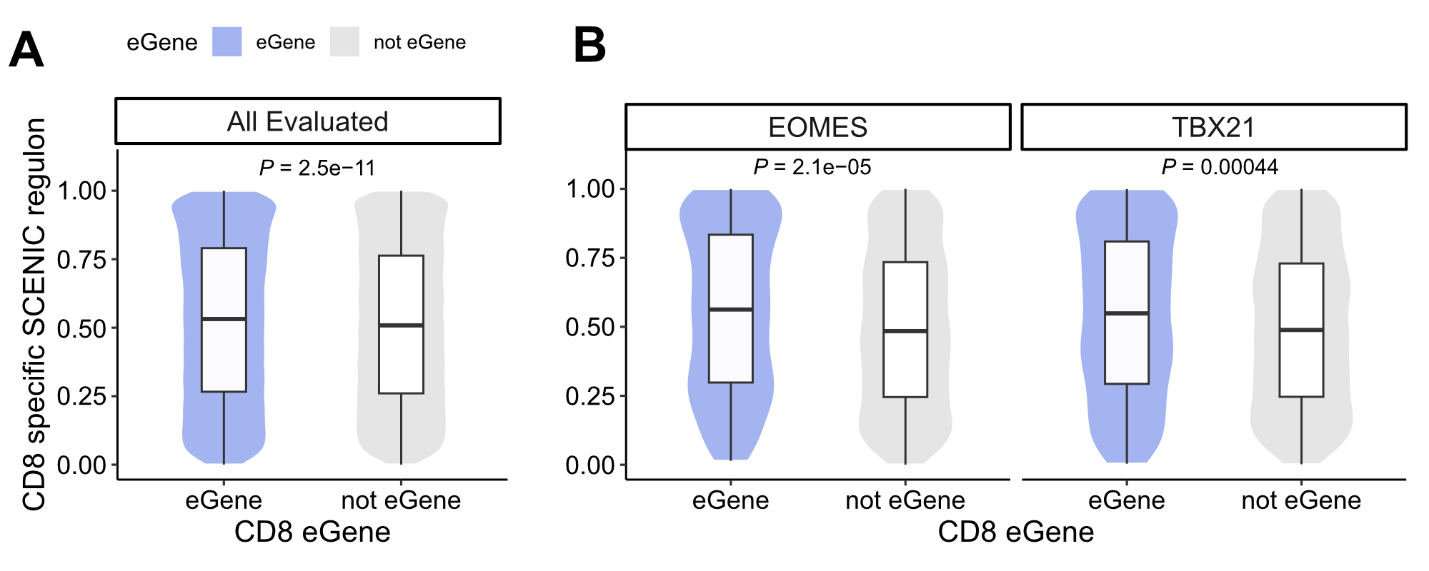

**Supplementary Figure S8. Comparison of importance rank from CD8 T-associated regulon by eGene.** **A** Combined comparison of the rank of importance metric in CD8 T-specific regulon based on eGene status. **B** Comparisons of TBX21, EOMES regulon in CD8 T cells based on eGene status.
